# Molecular evolution and characterization of domestic duck (*Anas platyrynchos*) and Goose (*Anser indicus*) with reference to its wild relatives through whole mitochondrial genome sequencing

**DOI:** 10.1101/2022.07.19.500621

**Authors:** Aruna Pal, Manti Debnath, Argha Chakraborty, Samiddha Banerjee, Abantika Pal

**Affiliations:** West Bengal University of Animal and Fishery Sciences, 37, K.B.Sarani, Kolkata-37

## Abstract

It is important to study the evolution and domestication of the domesticated duck (*Anas platyrynchos*) population from the wide range of wild relatives of Anas spp. Whole mitochondrial genome sequencing was attempted for Anas platyrynchos (Bengal duck) and Anser indicus (goose) from same geographical region. The study deals with the Molecular evolution of domestic duck based on mitochondrial gene due to its sequence variability, and to find out the phylogenetic relationships among *Anas platyrynchos* and its wild relatives. In this study we have used 45 wild species of Anas spp to study the mitochondrial genes and phylogenomics. Our result signifies that duck species were effectively discriminated with respect to mitochondrial genes, which could then be used for an appropriate genetic conservation program for the wild duck and domestic duck breeds. The DNA sequences from any unknown sample of the mitochondrial gene may be determined and can compare with those on a DNA database and can do blast for phylogenetic analysis of unknown wild duck, which gives its future scope. In silico analysis for 3D structure for *Anas platyrynchos* with the closest relative as *Anas poecilorhyncha* (Indian spot-billed duck) was attempted. *Anas platyrynchos* was also compared with *Anser indicus*. Therefore, this experiment was conducted to investigate the genetic diversity of West Bengal wild ducks with reference to its wild relatives based on mitochondrial gene.

## Introduction

Domestic duck has an important role in provision of animal protein and is reported to have a high genetic diversity, compared to other populations^**1**^. Ducks may be regarded as the second most important poultry species reared for egg and meat. Indigenous duck has been regarded as a hardy breed^**2**^ with better disease resistance ability^**3**^. In our lab, we have characterized certainfunction genes for duck^**4–13**^.

Whole mitochondrial genome for Anas platyrynchos was mapped as circular DNA with 37 genes without any intronic region^**14**^. They comprise of 13 polypeptide genes, D loop and RNA genes. It is interesting to note that some of the genes are negatively regulated^**14**^. One of the striking challenges of modern biology is to develop precise and dependable technologies for a speedy screening of DNA sequence dissimilarity. The quantity of mitochondria varies significantly between different organisms and even in between tissues also within the same organism^**15**^. Molecular classification is essential for the diagnosis, treatment and for the control of infections caused by different pathogens. In modern years, a variety of DNA-based approaches have been developed for the identification or classification of individuals for taxonomic groups.

The biodiversity and intra-specific genetic diversity are interrelated with each other and determine the promising community to stay alive and grow^**16**^. The molecular markers as well as DNA sequencing used as good markers for taxonomy and phylogenetic relationships among species analysis. On the other way, for phylogenetic tree mitochondrial DNA (mtDNA) sequences is more useful because the mtDNA is independent and more conservative as well as simpler than genomic DNA & able to give information about maternal inheritance^**17**^.

Researches indicate that proved that among mitochondrial genes, cytochrome b (mt Cytb) gene is efficient tool with high power of inequity for species identification as well as for characterization in both the cases like taxonomy and forensic science^**18,19**^. Study the rate of base substitution on mtDNA is 5-10 times if we compared it with any other nuclear gene of a wide variety of birds, which are closely interrelated with those species belong to same family or genera^**20**^. In Indonesia Purwantini and workers have identified the genetic diversity of Indonesian local duck population by PCR-RFLP method; those were Magelang, Tegal, Mojosari, Bali and Alabio ducks based on polymorphism of mtDNA D-loop^**21,22**^. Genetic diversity is essential for duck breeding management as well as for the morphometric diversity. The genetic diversity and morphometric are crucial for the development of duck breeding program appropriate in each country^**23**^.

Species identification can be done by nucleotide sequence investigation of the cytochrome b (cytb) gene^**24**^. They identified different biological specimens from miscellaneous vertebrate animals by extracting & amplifying DNA of 44 different animal species covering the 5 major vertebrate groups for example amphibians, mammals, fishes, birds, and reptiles. The sequences were used to identify the biological origin by aligning to cytb gene sequence entries in nucleotide databases. They had submitted their sequences to the GenBank. The mitochondrial association with diseases have been detected through the hypermethylation of mitochondrial CYTB and COX II genes. Decreased mtDNA has an important role in Alzheimer’s disease pathology, which may unwrap a latest porthole for Alzheimer’s disease therapy. They performed their experiment in an APP/PS1 transgenic mouse model of Alzheimer’s disease using pyrosequencing of the mtDNA methylation changes of the CYTB and COX II genes^**25**^.

Therefore, this experiment was conducted to investigate the genetic diversity of west Bengal wild ducks with reference to its wild relatives based on mitochondrial gene as well as available sequences for other mitochondrial genes..

## Materials and Method

### Whole mitochondrial genome sequencing of Anas platyrynchos and Anser indicus

Whole mitochondrial genome analysis (WMGA) was conducted for Bengal duck from West Bengal of India (Genebank accession number MN011574). The mitochondrial map was generated. The steps for next generation sequencing studies (Whole mitochondrial genome sequencing) involves isolation of mtDNA, qualitative and quantitative analysis of g-DNA: PCR amplified with COX-2(mt specific), GAPDH and Beta actin primers to validate. The next step involves preparation of library: The paired-end sequencing library (NEBNext Ultra DNA Library Preparation Kit), quantity and quality check (QC) of library on Bioanalyzer: Bioanalyzer 2100 (Agilent Technologies) using High Sensitivity (HS) DNA chip. The final step is cluster Generation and Sequencing. The adapters are designed so as to allow selective cleavage.

We had retrieved the complete mitochondrial sequences for other Anas spp from gene bank for the sequences available. Whenever the complete mitochondrial sequences were not available, sequences were retrieved for selected mitochondrial genes from gene bank.

### Sequence alignment and phylogenetic analysis

Sequences were imported into nucleotide BLAST (http://blast.ncbi.nlm.nih.gov/Blast.cgi) to recall comparable sequences from NCBI GenBank.Comparative alignment and phylogenetic analyses were performed by the help of MAAFT software^**26**^.

### Three dimensional structure prediction and model quality assessment

The templates which possessed highest sequence identity with our target template were identified by using PSI-BLAST (http://blast.ncbi.nlm.nih.gov/Blast). The homology modelling was used to build Cytochrome B native 3D structure based on homologous template structures using PHYRE2 server^**27**^. Predicts the 3D structure of a protein sequence based on HMM-HMM alignment techniques. For a given sequence, it detects known homologs based on PSI-Blast, constructs a hidden Markov model (HMM) of the sequence based on the detected homologs and scans this HMM against a database of HMMs of known protein structures. The 3D structures were visualized by PyMOL (http://www.pymol.org/) which is an open source molecular visualization tool. Subsequently, the mutant model was generated using PyMoL tool. The Swiss PDB Viewer was employed for controlling energy minimization. SAVES (Structural Analysis and Verification Server), an integrated server (http://nihserver.mbi.ucla.edu/SAVES/) is used for the structural evaluation of predicted 3D model of derived polypeptide.It is also used for assessing the stereochemical quality assessment of 3D predicted model. The ProSA (Protein Structure Analysis) web server (https://prosa.services.came.sbg.ac.at/prosa) was used for refinement and validation of protein structure^**28**^. The ProSA was used for checking model structural quality with potential errors and the program shows a plot of its residue energies and *Z*-scores which determine the overall quality of a model. The solvent accessibility surface area of the Cytochrome B was generated by using NetSurfP server (http://www.cbs.dtu.dk/services/NetSurfP/)^**29**^. TM align software was utilized for alignment of the 3 D structures of different variants of Cytochrome B protein pertaining to Anas spp^**30**^ along with other related bioinformatics software^**31**^.

## Results

### Whole mitochondrial genome sequencing for Anas platyrynchos and Anser indicus

Whole mitochondrial genome sequencing has revealed 37 genes with 13 polypeptide sequences, one control region (D loop) and RNAs for both *Anas platyrynchos* (Bengal duck) genome (Gene bank accession number MN011574.1) and *Anser indicus* (Goose from West Bengal). The mitochondrial map for *Anas platyrynchos* (Bengal duck) is being depicted in Fig 1, whereas for *Anser indicus* (Goose from West Bengal) is represented in Fig 2.

**Figure 1:**
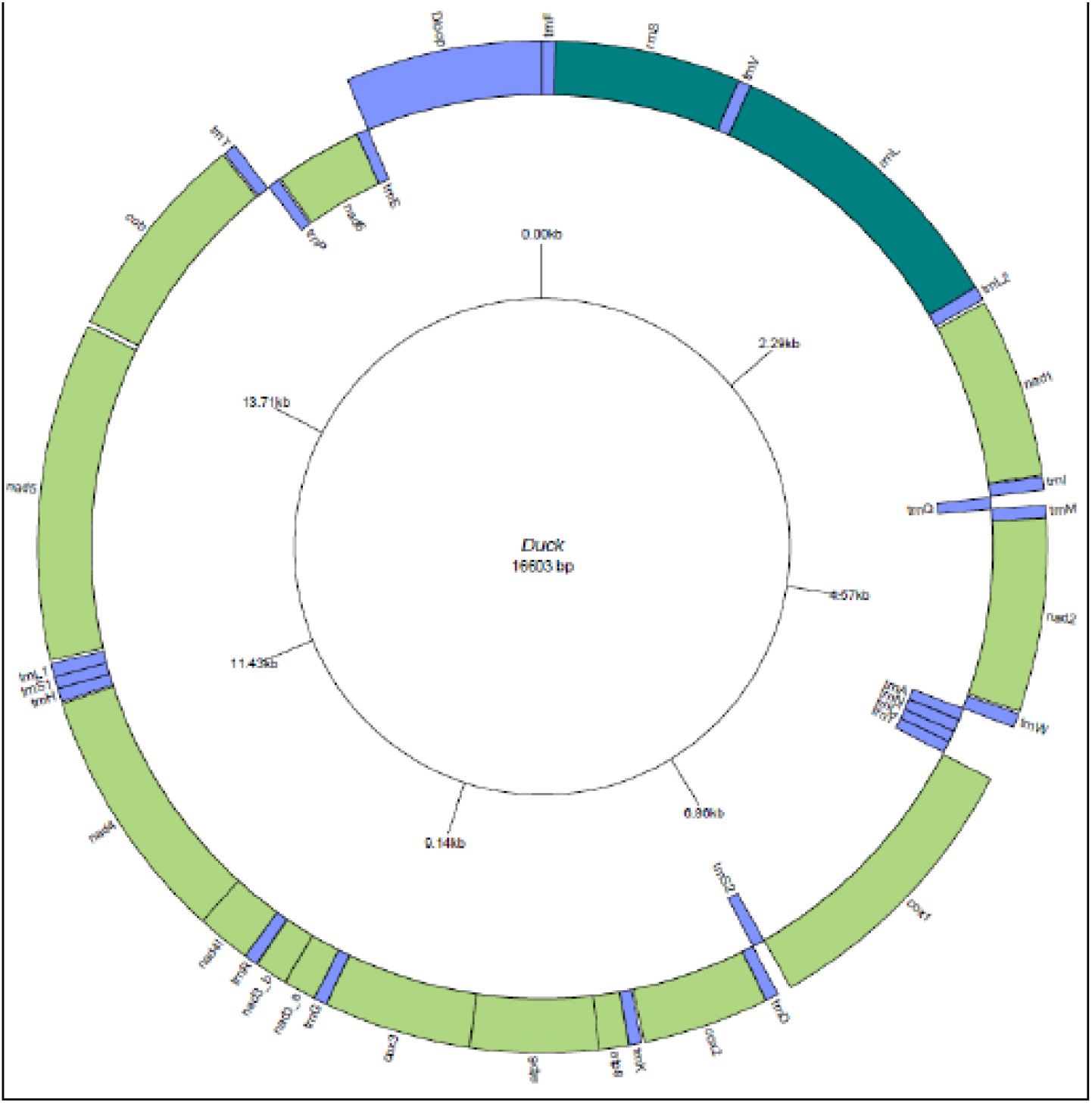
Circular map of Duck mitochondrial genome drawn using Genome Vx representing 37 protein coding genes with total genome size of 16,603 bp.

**Figure 2:**
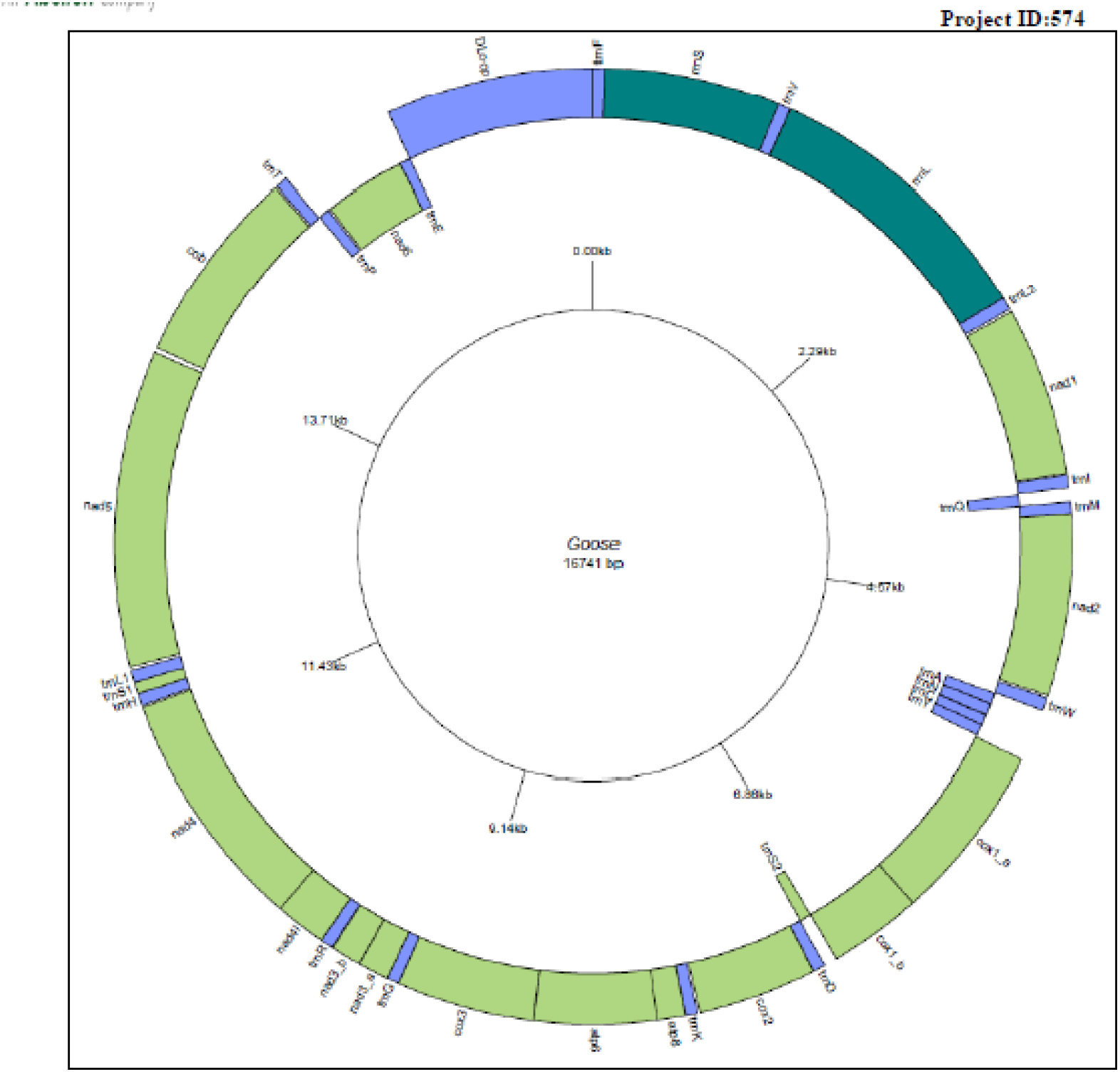
Circular map of Goose mitochondrial genome drawn using Genome Vs representing 37 protein coding genes with total genome size of 16,741 bp.

### Phylogenetic analysis of Anas platyrynchos with other Anas spp. Based on Cytochrome B genes

Phylogenetic tree for Cytochrome B gene of Anas platyrynchos with cytochrome B gene of other Anas spp (Fig 3).

**Fig 3:**
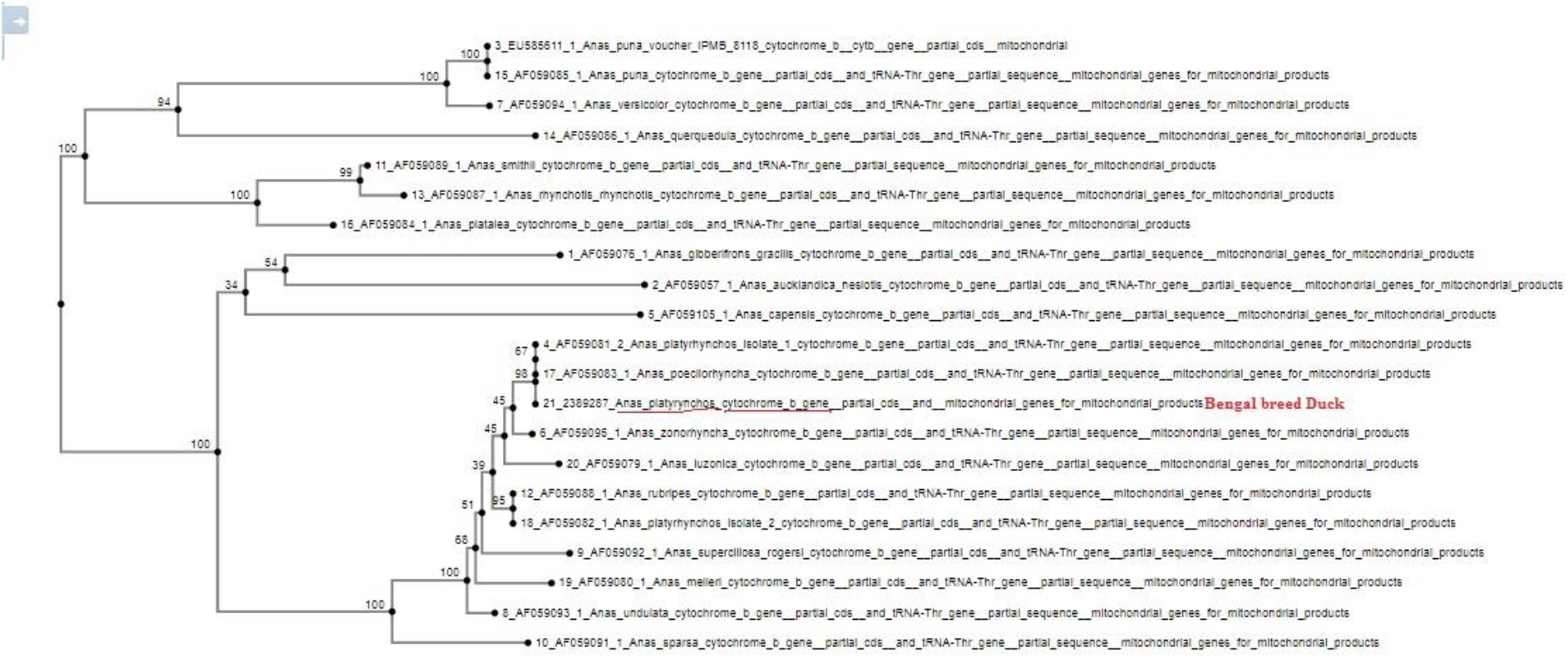
Phylogenetic analysis for *Anas platyrynchos* with other *Anas spp*.

### Comparison of cytochrome B in *Anas platyrynchos* and its close relative

The genetically closest relative for Bengal duck was observed to be Anas poecilorhyncha (Indian spot-billed duck), complete genome GenBank: KC466567.1 among other wild duck species. Percent identity matrix developed by Clustal 12.1 depicts 99.7 percent identity among these two species. Amino acid changes were not detected for these two species. These changes in amino acid has been reflected in 3D structure of cytochrome B of both the closest related species. The next closest species is Anas zonorhyncha mitochondrion, complete genome GenBank: MZ593724.1.

### Characterization and domain identification for Cytochrome B gene of Anas platyrynchos

Mitochondrial Cytochrome B has been characterized as 380 amino acids, ncbi gene bank protein id as QVX28267.1. It is a part of circular mitochondrial DNA. The sequence derived from Anas platyrynchos mitochondrion complete genome (Gene bank accession number MN011574.1).

### Comparison of 3D structural analysis for the derived peptides for mitochondrial gene

3D structural analysis for derieved peptide for thirteen polypeptide genes have been depicted for *Anas platyrynchos* with *Anas poecilorhyncha* and *Anser indicus*. 3D structure for *Anas platyrynchos* with *Anas poecilorhyncha* does not reveal any RMSD or any structural difference (Fig 4). However significant structural differences were observed for *Anas platyrynchos* with *Anser indicus*. Differences were observed for ND2 (RMSD:0.15), Cox2 (RMSD: 0.55), ND5 (RMSD: 0.44) ND6 (RMSD:2.68).

**Fig4:**
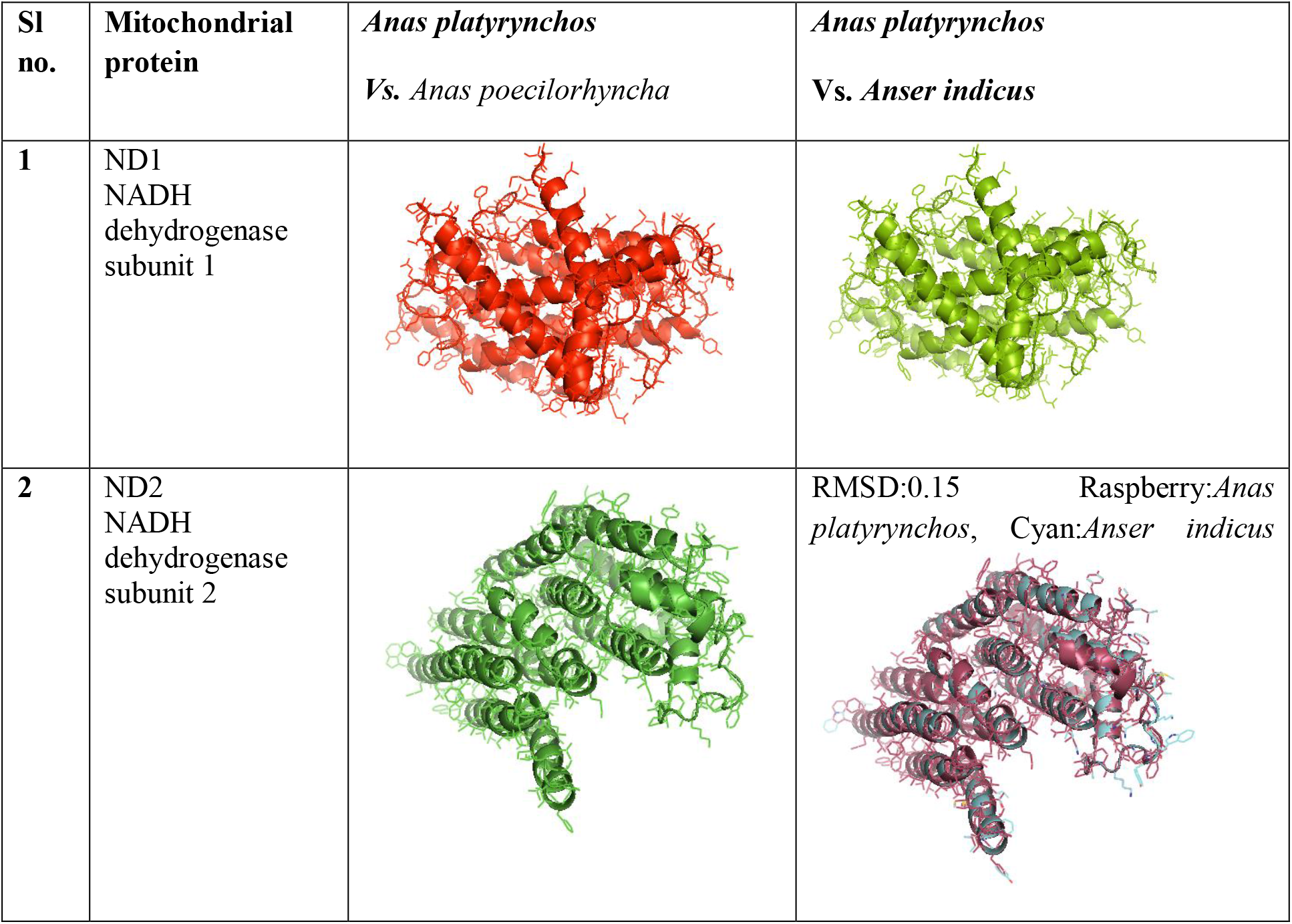
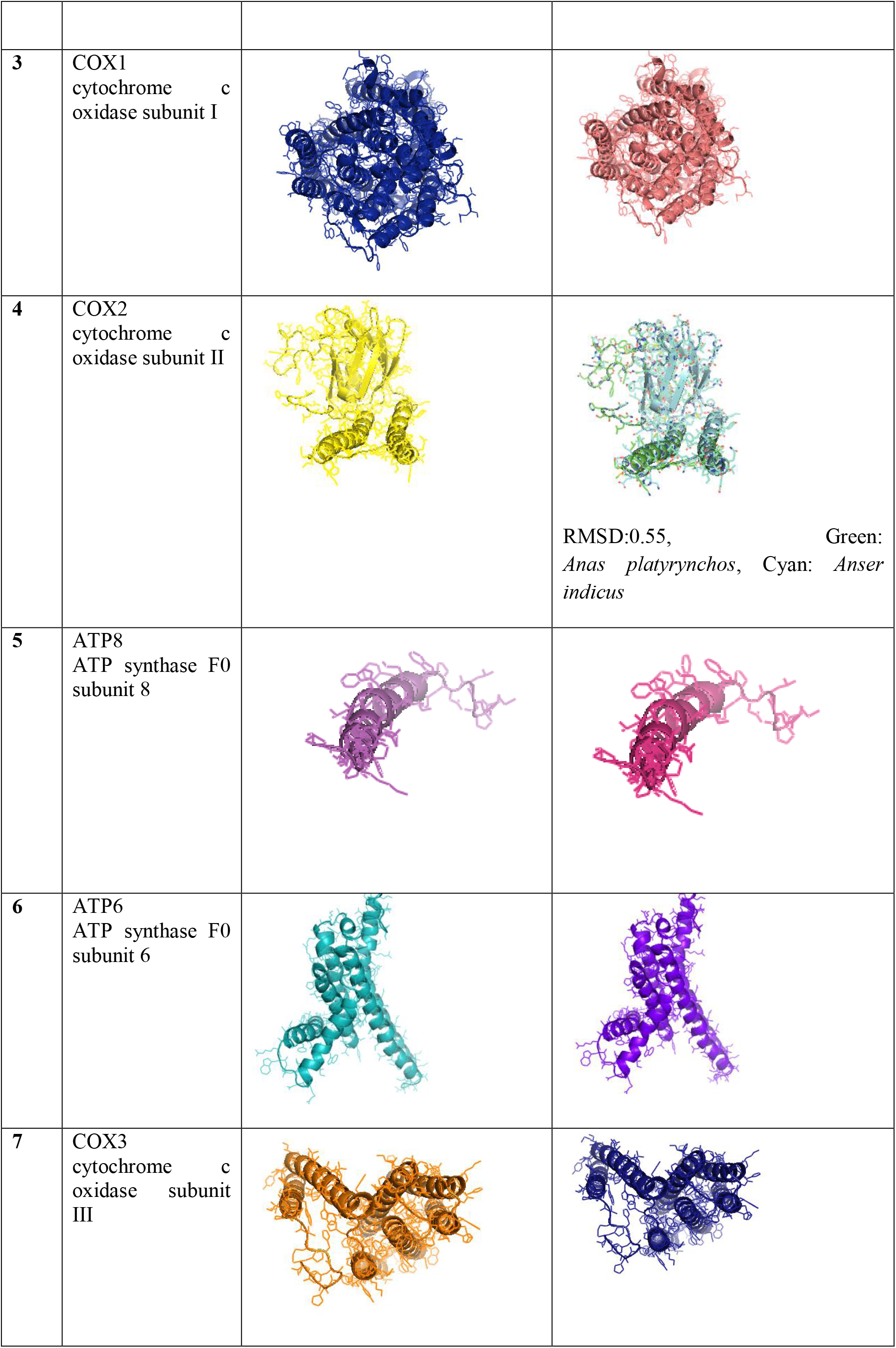
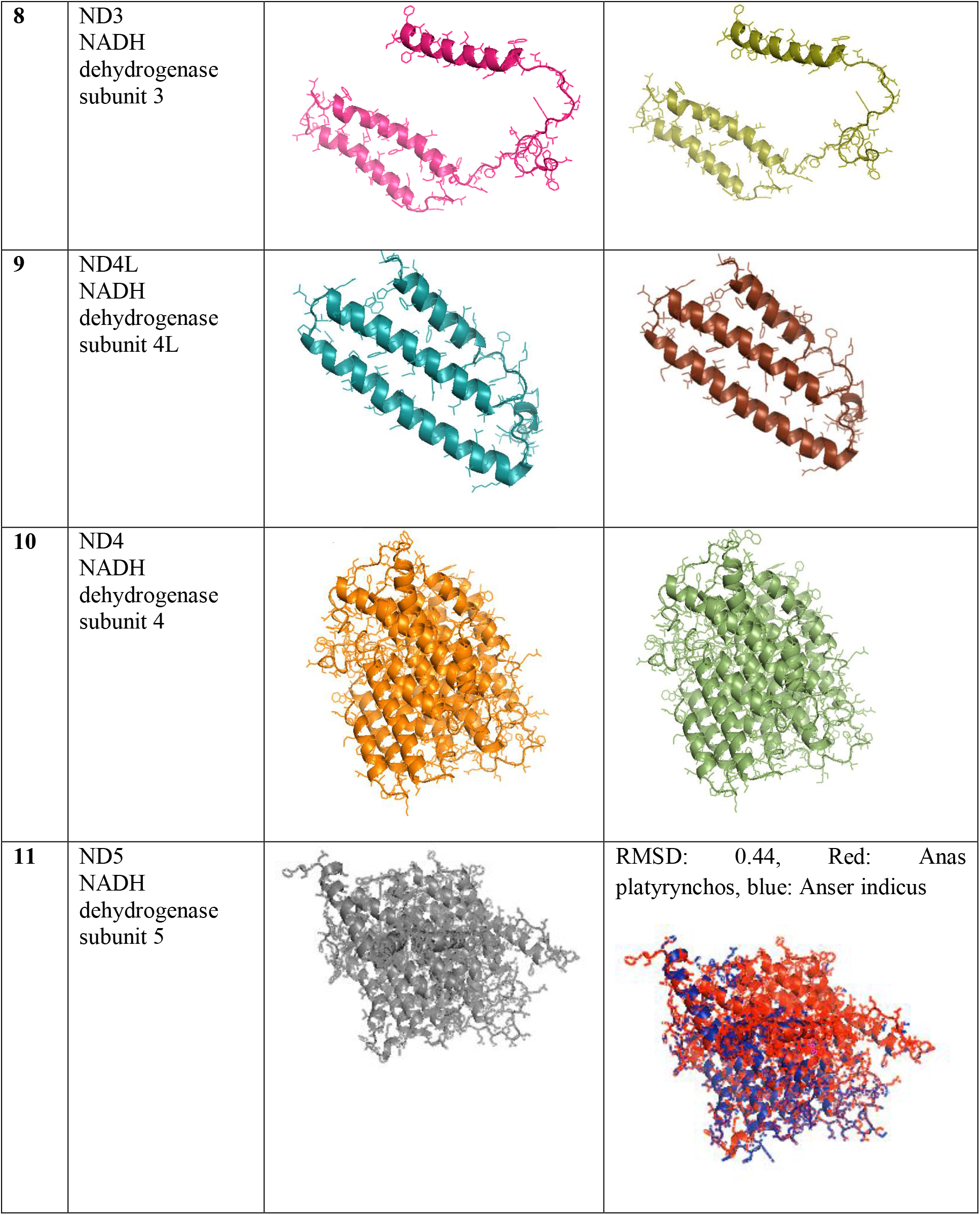
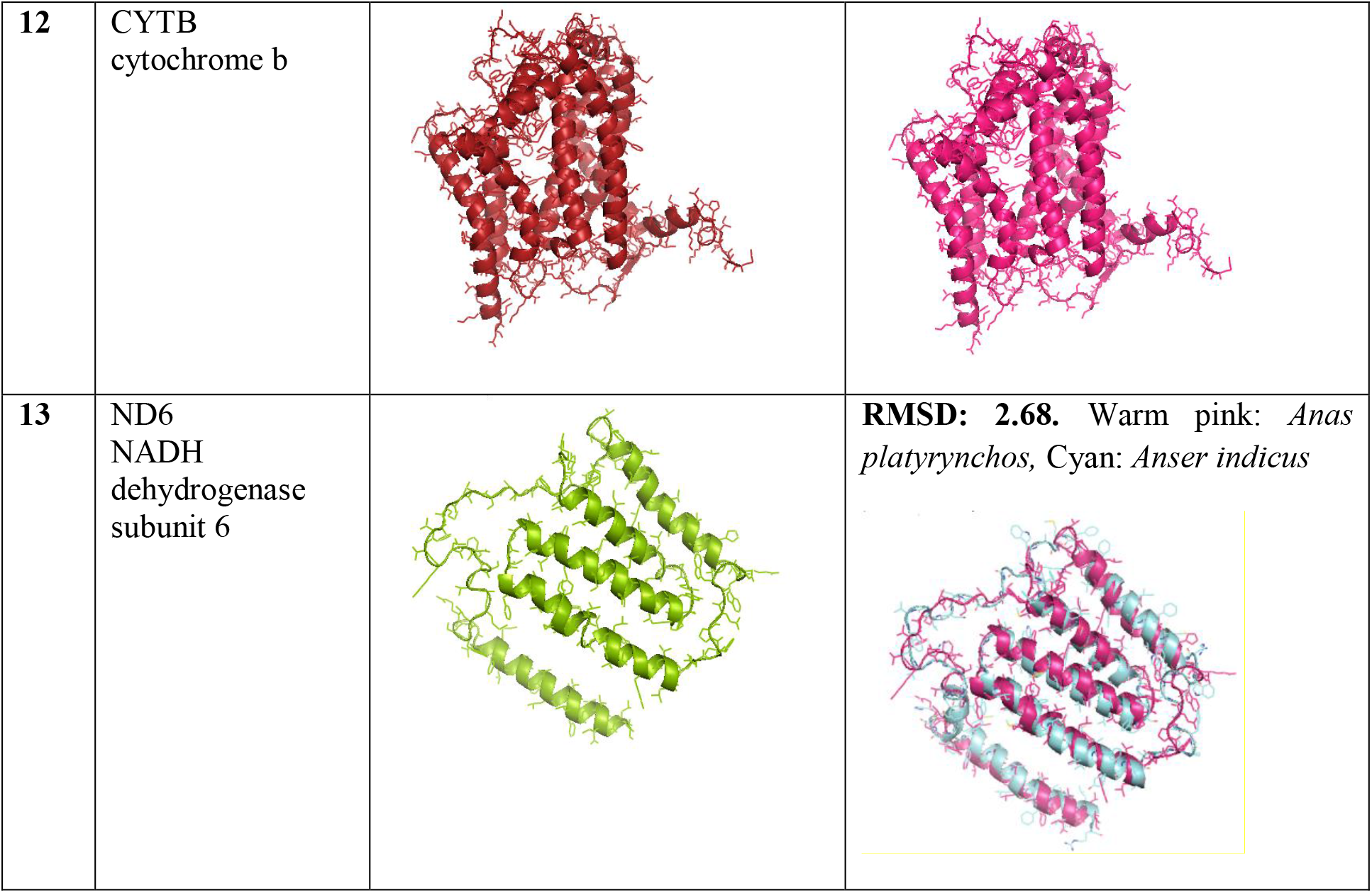
3D structural analysis for the derieved polypeptides from coding regions of mitochondrial genes:

### Comparison of ND6 sequence between Anas platyrynchos with Anser indicus

The variations in nucleotide sequences have been detected at amino acid position of Anas platyrynchos with Anser indicus as V13A, H90C, A92V, L94F, A95V, V97I, L99F, L104F, G112K, V116S, S120N, T125V, L128S, L136F, V141A, W149C (Fig 5)

**Fig 5:**
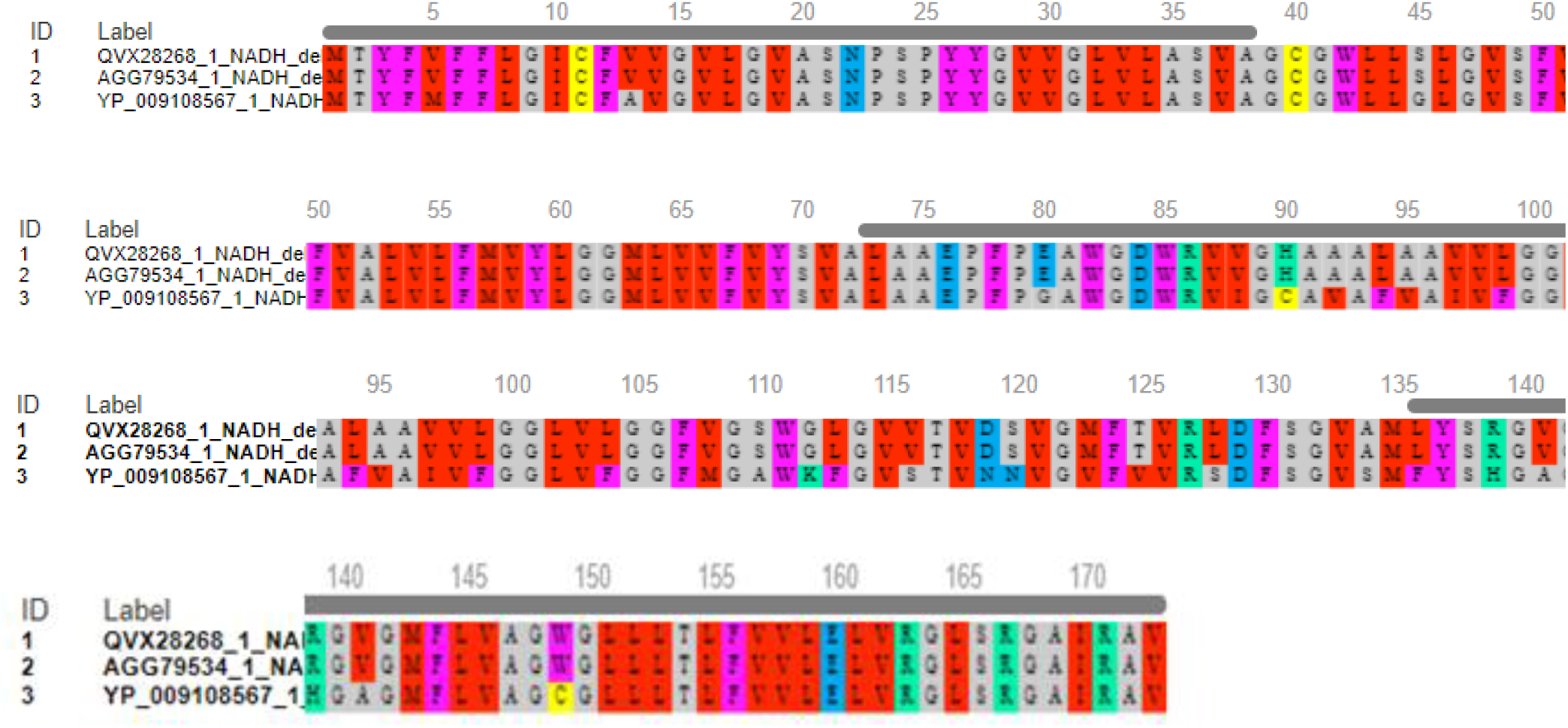
Amino acid alignment for *Anas platyrynchos, Anas poecilorhyncha and Anser indicus*

## Discussion

Current day domestic ducks (*Anas platyrynhos*) are evolved as a promising poultry species with some unique traits of highly nutritious egg, fetching higher price in the market. The most remarkable characteristics for domesticated duck is its ability to resist most of the common avian diseases. The evolution and domestication of Anas platyrynchos is important to study. Migratory birds of Anas spp and others are important for spread of certain diseases of extreme zoonotic importance as avian influenza. In the current study, we have studied the phylogenetic analysis of *Anas platyrynchos with other Anas species* and *Anser indicus*. We have retrieved 21 sequences for Anas spp under current study. *Anas platyrynchos* has been observed to be genetically closest to *Anas poecilorhyncha*. Next closest relative was observed to be *Anas zonorhyncha*. Similar study was reported earlier based on Cytochrome B and ND2 for phylogenetic analysis of Anseriformes^**32**^. They have studied 45 spp of Anseriformes comprising of both Anatinae and Anserinae.

In the current study, we have studied whole mitochondrial genome analysis comprising of 13 protein coding genes, 2 rRNA genes (12S rRNA and 16S rRNA), 22 tRNA genes, and 1 control region (CR) each for Anas platyrynchos, *and Anser indicus* and retrieved complete mitochondrial genome sequence for *Anas poecilorhyncha* from gene bank. Comparison of derived amino acids have revealed certain non-synonymous mutations leading to amino acid variations for *Anas platyrynchos* and *Anser indicus*. We could identify V13A, H90C, A92V, L94F, A95V, V97I, L99F, L104F, G112K, V116S, S120N, T125V, L128S, L136F, V141A, W149C amino acid differences.

Protein sequence differences as well as structural differences in derived 3D structures for all the polypeptide coding genes have been studied for *Anas platyrynchos and Anser indicus*. Structural similarity has been observed for derived 3D structure for mitochondrial protein as ND1, ND3, Cox1, ATP6, ATP8, Cox3, ND3, ND4, ND4L, CytB. Differences were observed for ND2 (RMSD:0.15), Cox2 (RMSD: 0.55), ND5 (RMSD: 0.44), ND6 (RMSD:2.68). These studies for evolution based on mitochondrial protein structural analysis is unique in nature and studied for the first time to detect evolutionary relationship among *Anas platyrynchos and Anser indicus*. In one of our earlier studies, we have analyzed molecular studies for Anas platyrynchos sequences across the globe, based on derived protein structure for all the mitochondrial genes, thirteen in number.

Earlier we have already studied the functional association of cytochrome B gene with health in sheep model^**33**^ and reproduction in pig model^**34**^. We have studied in detail the bioinformatics analysis for Cytochrome B gene with respect to its functional association. We have also studied the phylogenetic studies of sheep from West Bengal based on Cytochrome B gene^**35**^. We have equally studied the phylogenetic relationship among the duck population from different agroclimatic conditions based on biomorphometric characteristics^**36**^. We have also studied the remdies for analysis of mitochondrial defects ^**37**^.Certain reports are available for phylogenetic studies of duck with respect to individual mitochondrial genes^**38–47**^, but whole mitochondrial genome sequencing employed for studying molecular phylogeny for duck and related species is unique in nature.

## Acknowledgement

The authors are thankful to Department of Biotechnology, Ministry of Science and Technology, Govt. of India (Grant number BT/PR24310/NER/95/649/2017) and Science and engineering research Board, Department of Science and Technology, Govt. of India (Grant no. EMR/2016/003554) for providing the financial support. The technical and financial support by Vice-Chancellor, West Bengal University of Animal and Fishery Sciences is duly acknowledged. Thanks to Director, AH & VS, Animal Resource Development Department, Govt. of West Bengal.

## Notes

### Competing Interest Statement

The authors have declared no competing interest.

